# The Image Data Resource: A Scalable Platform for Biological Image Data Access, Integration, and Dissemination

**DOI:** 10.1101/089359

**Authors:** Eleanor Williams, Josh Moore, Simon W. Li, Gabriella Rustici, Aleksandra Tarkowska, Anatole Chessel, Simone Leo, Bálint Antal, Richard K. Ferguson, Ugis Sarkans, Alvis Brazma, Rafael E. Carazo Salas, Jason R. Swedlow

**Affiliations:** Centre for Gene Regulation & Expression & Division of Computational Biology, University of Dundee, Dundee, Scotland, UK; European Molecular Biology Laboratory, European Bioinformatics Institute, Hinxton, United Kingdom; Pharmacology & Genetics Departments and Cambridge Systems Biology Centre, University of Cambridge, Cambridge, UK; School of Cell and Molecular Medicine, University of Bristol, Bristol, UK; Center for Advanced Studies, Research, and Development in Sardinia (CRS4), Pula(CA), Italy; LOB, Ecole Polytechnique, CNRS, INSERM, Université Paris-Saclay, Palaiseau, France

## Abstract

Access to primary research data is vital for the advancement of science. To extend the data types supported by community repositories, we built a prototype Image Data Resource (IDR) that collects and integrates imaging data acquired across many different imaging modalities. IDR links high-content screening, super-resolution microscopy, time-lapse and digital pathology imaging experiments to public genetic or chemical databases, and to cell and tissue phenotypes expressed using controlled ontologies. Using this integration, IDR facilitates the analysis of gene networks and reveals functional interactions that are inaccessible to individual studies. To enable re-analysis, we also established a computational resource based on IPython notebooks that allows remote access to the entire IDR. IDR is also an open source platform that others can use to publish their own image data. Thus IDR provides both a novel on-line resource and a software infrastructure that promotes and extends publication and re-analysis of scientific image data.

---

Much of the published research in the life sciences is based on image datasets that sample 3D space, time, and the spectral characteristics of detected signal (e.g., photons, electrons, proton relaxation, etc) to provide quantitative measures of cell, tissue and organismal processes and structures. The sheer size of biological image datasets makes data submission, handling and publication extremely challenging. An image-based genome-wide “high-content” screen (HCS) may contain over a million images, and new “virtual slide” and “light sheet” tissue imaging technologies generate individual images that contain gigapixels of data showing tissues or whole organisms at subcellular resolutions. At the same time, published versions of image data often are mere illustrations: they are presented in processed, compressed formats that cannot convey the measurements and multiple dimensions contained in the original image data and that can no longer be easily subjected to re-analysis. Furthermore, conventional publications neither include the metadata that define imaging protocols, biological systems and perturbations nor the processing and analytic outputs that convert the image data into quantitative measurements.

Several public image databases have appeared over the last few years. These provide online access to image data, enable browsing and visualisation, and in some cases include experimental metadata. The Allen Brain Atlas, the Human Protein Atlas, and the Edinburgh Mouse Atlas all synthesise measurements of gene expression, protein localization and/or other analytic metadata with coordinate systems that place biomolecular localisation and concentration into a spatial and biological context ^1–3^. Similarly, many other examples of dedicated databases for specific imaging projects exist, each tailored to its aims and its target community ^4–8^. There are also a number of public resources that serve as true scientific, structured repositories for image data, i.e., that collect, store and provide persistent identifiers for long-term access to submitted datasets, as well as provide rich functionalities for browsing, search and query. One archetype is the EMDataBank, the definitive community repository for molecular reconstructions recorded by electron microscopy ^9^. *The Journal of Cell Biology* has built the JCB DataViewer, which publishes image datasets associated with its on-line publications ^10^. The CELL Image Library publishes several thousand community-submitted images, some of which are linked to publications ^11^. FigShare stores 2D pictures derived from image datasets, and can provide links for download of image datasets (http://figshare.com). The EMDataBank recently has released a prototype repository for 3D tomograms, the EMPIAR resource ^12^. Finally, the BioStudies and Dryad archives include support for browsing and downloading image data files linked to studies or publications ^13^ (https://datadryad.org/). Some of the these provide a resource for a specific imaging domain (e.g., EMDataBank) or experiment (e.g., Mitocheck), while others archive datasets and provide links to a related publication available at an external journal’s website (e.g., BioStudies). However, no existing resource links several independent biological imaging datasets to provide an “added value” platform, like the Expression Atlas achieves for a broad set of gene expression datasets ^14^ and UniProt delivers for protein sequence and function datasets ^15^.

Inspired by these “added value” resources, we have built a next-generation <u>I</u>mage <u>D</u>ata <u>R</u>esource (IDR) – an added value platform that combines data from multiple independent imaging experiments and from many different imaging modalities, integrates them into a single resource, and makes the data available for re-analysis in a convenient, scalable form. IDR provides, for the first time, a prototyped resource that supports browsing, search, visualisation and computational processing within and across datasets acquired from a wide variety of imaging domains. For each study, metadata related to the experimental design and execution, the acquisition of the image data, and downstream interpretation and analysis are stored in IDR alongside the image data and made available for search and query through a web interface and a single API. Wherever possible, we have mapped the phenotypes determined by dataset authors to a common ontology. For several studies, we have calculated comprehensive sets of image features that can be used by others for reanalysis and the development of phenotypic classifiers. By harmonising the data from multiple imaging studies into a single system, IDR users can query across studies and identify phenotypic links between different experiments and perturbations.

## Results

### Current IDR

IDR is currently populated with 24 imaging studies comprising 35 screens or imaging experiments from the biological imaging community, most of which are related to and linked to published works (Table 1). IDR holds ∼42 TB of image data in ∼36M image planes and ∼1M individual experiments, and includes all associated experimental (e.g., genes, RNAi, chemistry, geographic location), analytic (e.g., submitter-calculated regions and features), and functional annotations. Datasets in human cells (e.g., http://idr-demo.openmicroscopy.org/webclient/?show=well-45407; http://idr-demo.openmicroscopy.org/webclient/?show=well-547609) and fungi (e.g., http://idr-demo.openmicroscopy.org/webclient/?show=well-590686; http://idr-demo.openmicroscopy.org/webclient/?show=well-469267),super resolution 3DSIM images of centrosomes (http://idr-demo.openmicroscopy.org/webclient/?show=dataset-51) and dSTORM images of nuclear pores (http://idr-demo.openmicroscopy.org/webclient/?show=dataset-61), a comprehensive chemical screen in human cells (http://idr-demo.openmicroscopy.org/webclient/?show=plate-4101), a live cell screen in human cells (Mitocheck; http://idr-demo.openmicroscopy.org/webclient/?show=well-771034) and histopathology whole slide images of tissues from several mouse mutants (http://idr-demo.openmicroscopy.org/webclient/?show=dataset-369) are included. Finally, imaging from Tara Oceans, a global survey of plankton and other marine organisms, is also included (http://idr-demo.openmicroscopy.org/webclient/?show=plate-4751). The current collection of datasets samples a variety of biomedically-relevant biological processes like cell shape, division and adhesion, at scales ranging from nanometre-scale localisation of proteins in cells to millimetre-scale structure of tissues from animals.

**Table 1.**
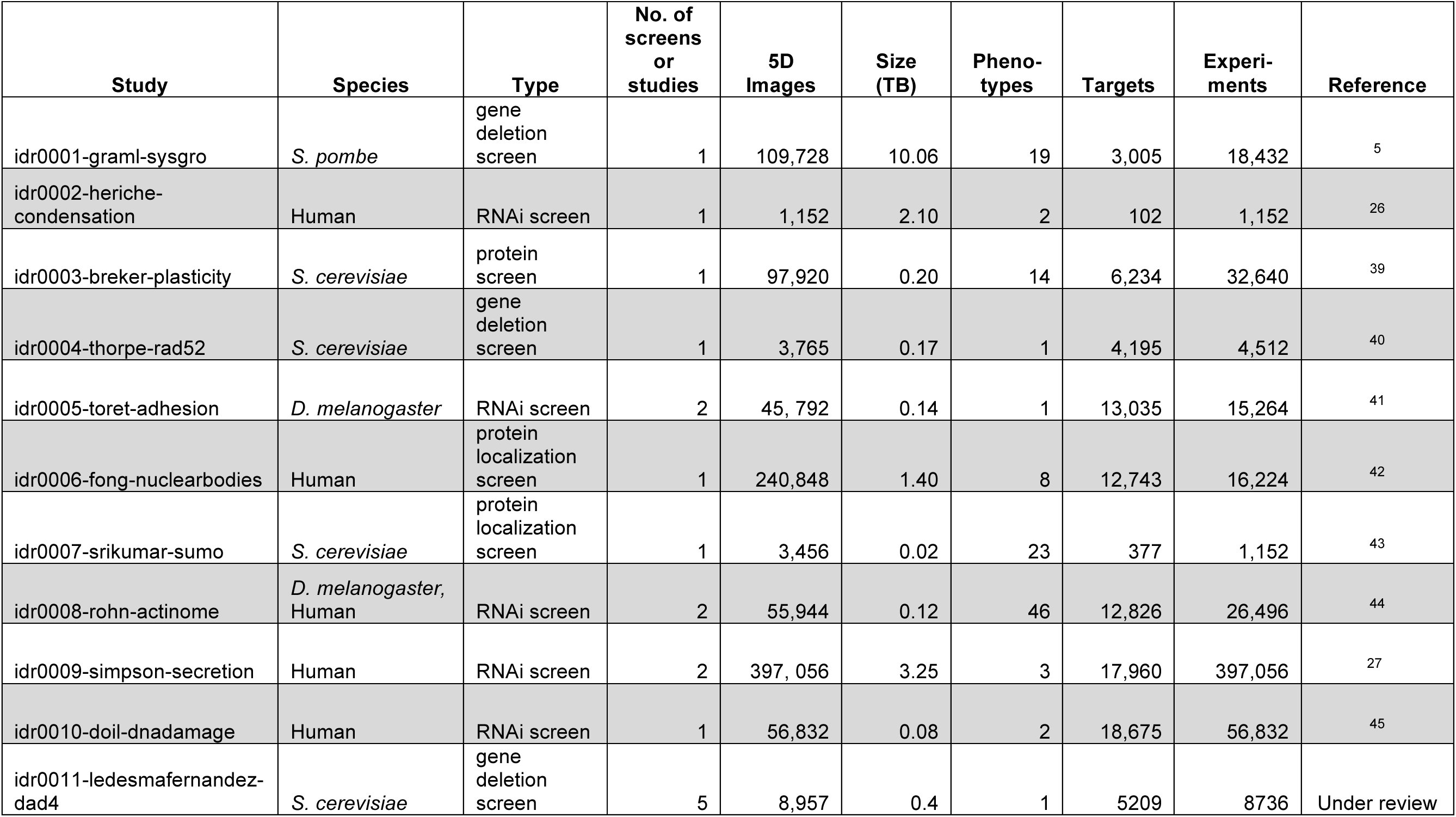
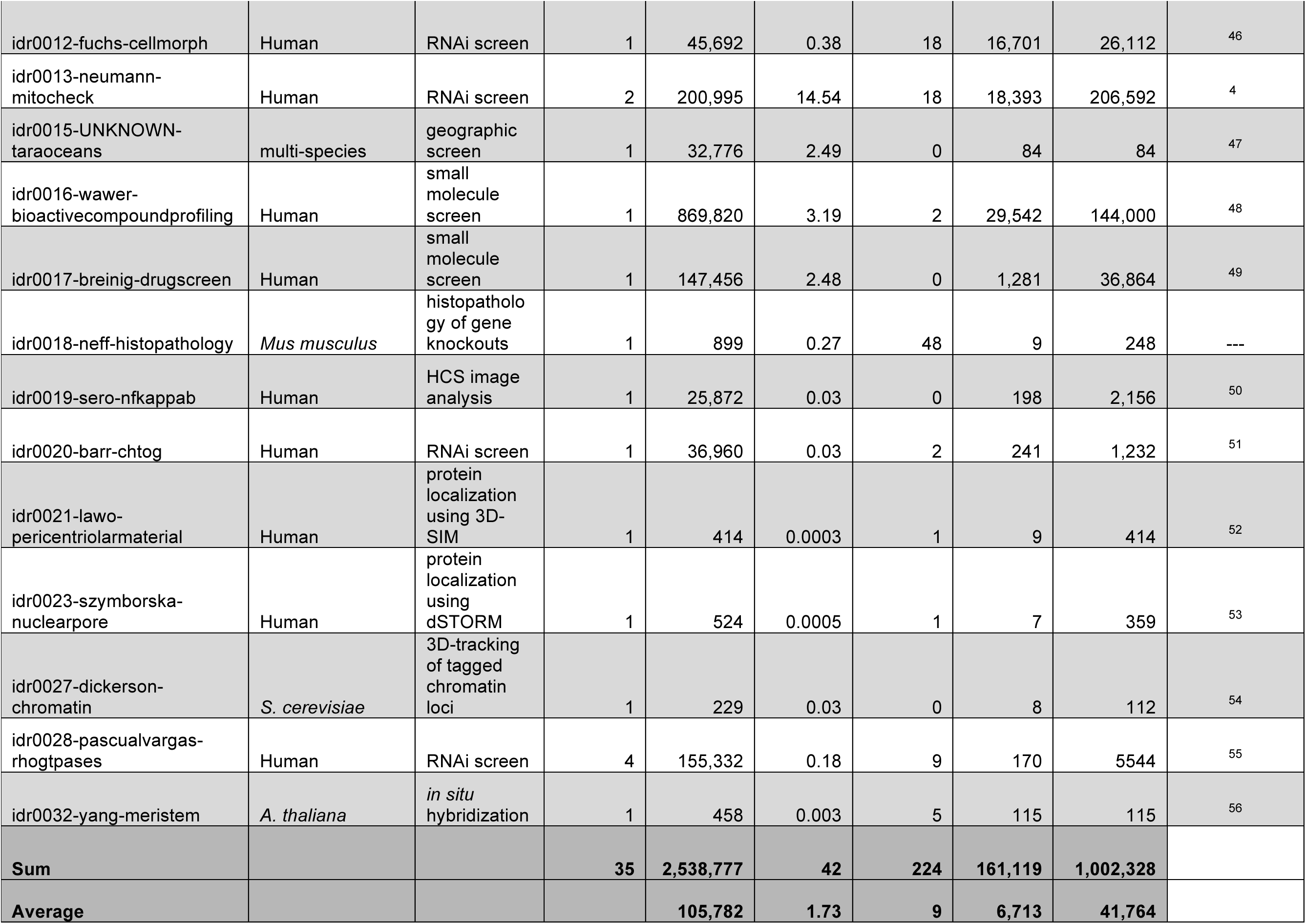
List of Datasets in IDR. The phenotype column contains the number of submitted phenotypes. The number of genes, compounds or proteins identified as targets for analysis is listed in the Targets column and the ‘Experiments’ column lists the number of individual wells in HCS studies or imaging experiments in non-screen datasets

### Genetic, Chemical and Functional Annotation in IDR

To enable querying across the different datasets stored in IDR, we have included annotations describing experimental perturbations (genetic mutants, siRNA targets and reagents, expressed proteins, cell lines, drugs, etc.) and phenotypes declared by the study authors either from quantitative analysis or visual inspection of the image data. Wherever possible, experimental metadata in IDR link to external resources that are the authoritative resource for those metadata (Ensembl, NCBI, PubChem, etc).

The result is that IDR is a sampling of phenotypes related to experimental perturbations across several independent studies. Many of the studies in IDR perturb gene function by mutation or siRNA depletion. To calculate the sampling of gene orthologues, we used Ensembl’s BioMart resource ^16^ to access a normalised list of gene orthologues. Overall, 19,601 gene orthologues are sampled, and 84.1% of gene orthologues are sampled more than 20 times. 90.3% of gene orthologues are sampled in three or more studies, so even in this early incarnation the phenotypes of perturbations in the majority of known genes are sampled in several different assays and organisms.

We also sought to normalise the phenotypes defined by submitting study authors in IDR. Functional annotations (e.g., “increased peripheral actin”) have been converted to defined terms in the Cellular Microscopy Phenotype Ontology (CMPO) or other ontologies ^17^, in collaboration with the data submitters (e.g., http://idr-demo.openmicroscopy.org/webclient/?show=image-109846). Overall, 88% of the functional annotations have links to defined, published controlled vocabularies. 158 different ontology-normalised phenotypes (e.g., “increased number of actin filaments”, “mitosis arrested”) are included in IDR, and 136 are reported by authors in only one study. Nonetheless, these phenotypes are well-sampled--the mean number of samples per phenotype, across HCS and other imaging datasets is 698 and the median is 144. This skewing occurs because some phenotypes are very common or are over-represented in specific assays, e.g. “protein localized in cytosol phenotype”, (CMPO_0000393; http://idr-demo.openmicroscopy.org/mapr/phenotype/?value=CMPO_0000393). Nonetheless, there are several cases where phenotypes are observed in multiple orthogonal assays. Two examples are the “round cell” phenotype (CMPO_0000118; http://idr-demo.openmicroscopy.org/mapr/phenotype/?value=CMPO_0000118) and the “increased nuclear size” phenotype (CMPO_0000140; http://idr-demo.openmicroscopy.org/mapr/phenotype/?value=CMPO_0000140). Figure 1 summarises the sampling of phenotypes across the current IDR datasets. Several classes of phenotypes are included, and many cases are sampled in thousands of individual experiments. In total, IDR includes more than one million individual experiments (Table 1), with ∼9 % annotated with experimentally observed phenotypes.

**Figure 1.**
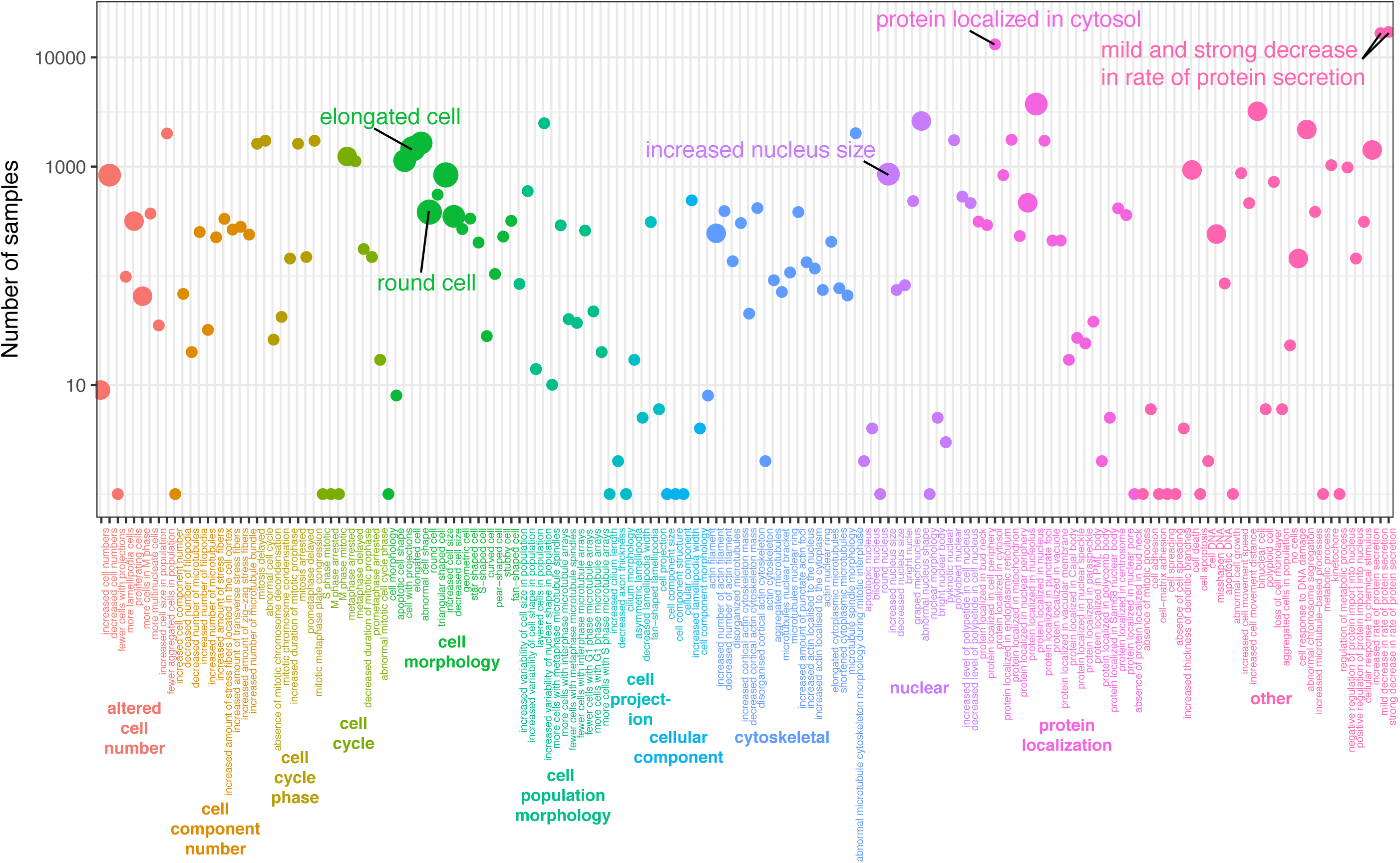
Sampling of Phenotypes in the IDR. The numbers of samples per phenotype. Each sample represents a well from a micro-well plate in a screen or image from a dataset. Wells annotated as controls were not included. User submitted phenotype terms were mapped to the CMPO terms shown here. Colours represent higher-level groupings of phenotype terms. Point size shows the number of studies (group of related screens) each phenotype is linked to with small, medium and large points representing 1, 2 or 3 studies respectively.

### Data Visualization in IDR

IDR integrates image data and metadata from several studies into a single resource. The current IDR web user interface (WUI) is based on the open source OMERO.web application ^18^ supplemented with a plugin allowing datasets to be viewed by ‘Study’, ‘Genes’, ‘Phenotypes’, ‘siRNAs’, ‘Antibodies’, ‘Compounds’, and ‘Organisms’ (see Supplemental Information). Using this architecture makes the integrated data resource available for access and re-use in several ways. Image data are viewable as thumbnails for each study (e.g., http://idr-demo.openmicroscopy.org/webclient/?show=plate-4349) and multi-dimensional images can be viewed and browsed (e.g., http://idr-demo.openmicroscopy.org/webclient/?show=well-45501 and http://idr-demo.openmicroscopy.org/webclient/?show=well-93714). Tiled whole slide images used in histopathology are also supported (e.g., http://idr-demo.openmicroscopy.org/webclient/?show=image-1920135). Where identified regions of interest (ROIs) have been submitted with the image data, these have been included and linked, and where possible, made available through the IDR WUI (e.g., http://idr-demo.openmicroscopy.org/webclient/?show=well-590686 and http://idr-demo.openmicroscopy.org/webclient/imgdetail/1230005/). IDR images, thumbnails and metadata are accessible through the IDR WUI and web-based API in JSON format (see Supplemental Information). They also can be embedded into other pages using the OMERO.web gateway (e.g., https://www.eurobioimaging-interim.eu/image-data-repository.html).

### Standardised Interfaces for Imaging Metadata

IDR integrates imaging data from many different, independent studies. These data were acquired using several different imaging modalities, in the absence of any over-arching standards for experimental, imaging or analytic metadata. While efforts like MIACA (http://miaca.sourceforge.net/), NeuroVault ^19^ MULTIMOT ^20^ and several other projects have proposed data standards in specific imaging subdomains, there is not yet a metadata standard that crosses all of the imaging domains potentially served by IDR. We therefore sought to adopt lightweight methods from other communities that have had broad acceptance ^21^ and converted metadata submitted in custom formats – spreadsheets, PDFs, MySQL databases, and Microsoft Word documents -- into a consistent tabular format inspired by the MAGE-TAB and ISA-TAB specifications ^22, 23^ that could then be used for importing semi-structured metadata like gene and ontology identifiers into OMERO. We also used the Bio-Formats software library to identify and convert well-defined, semantically-typed elements that describe the imaging metadata (e.g., image pixel size) as specified in the OME Data Model ^24, 25^. The resulting translation scripts were used to integrate datasets from multiple distinct studies and imaging modalities into a single resource. The scripts are publicly available (see Methods) and thus comprise a framework for recognising and reading a range of metadata types across several imaging domains into a common, open specification.

### Added Value of IDR

Because IDR links gene names and phenotypes, query results that combine genes and phenotypes across multiple studies are possible through simple text-based search. Searching for the gene SGOL1 (http://idr-demo.openmicroscopy.org/mapr/gene/?value=SGOL1) returns a range of phenotypes from four separate studies associated with mitotic defects (for example, CMPO_0000118, CMPO_0000305, CMPO_0000212, CMPO_0000344, etc.) ^4, 26^ but also an accelerated secretion phenotype (CMPO_0000246) in a screen for defects in protein secretion ^27^. A second example is provided in a histopathology study of tissue phenotypes in a series of mouse mutants. Knockout of carbonic anhydrase 4 (CAR4; http://idr-demo.openmicroscopy.org/mapr/gene/?value=Car4) in mouse results in a range of defects in homeostasis in the brain, rib growth and male fertility ^28–30^. Data held in IDR show abnormal nuclear phenotypes in several tissues from CAR4^-/-^ mice (e.g., GI: http://idr-demo.openmicroscopy.org/webclient/?show=dataset-153; liver: http://idr-demo.openmicroscopy.org/webclient/?show=image-1918940; male reproductive tract: http://idr-demo.openmicroscopy.org/webclient/?show=image-1918953). The human orthologue, CA4, is involved in certain forms of retinitis pigmentosa ^31, 32^. Data presented in IDR from the Mitocheck study show that siRNA depletion of CA4 in HeLa cells 4 also results in abnormally shaped nuclei (http://idr-demo.openmicroscopy.org/webclient/?show=well-828419) consistent with a defect in some aspect of the cell division cycle.

Phenotypes across distinct studies can also be used to build novel representations of gene networks. Figure 2A shows the gene network created when the gene knockouts or knockdowns that caused an elongated cell phenotype (CMPO_0000077) in studies in *S. pombe* and human cells are linked by queries to String DB and visualised in Cytoscape ^33^. The genes discovered in the three studies form interconnected, non-overlapping, complementary networks that connect specific macromolecular complexes to the elongated cell phenotype. For example, HELZ2, MED30, MED18 and MED20 are all part of the Mediator Complex, but were identified as “elongated cell” hits in separate studies using different biological models (idr0001-A, idr0008-B, idr0012-A, Figure 2B). In another example, POLR2G (from idr0012-A), PAF1 (from idr0001-A) and SUPT16H (from idr0008-B) were scored as elongated cell hits in these studies and are all part of the Elongation complex in the RNA Polymerase II transcription pathway. Finally, ASH2L (“elongated cell phenotype” in idr0012-A), associates with SETD1A and SETD1B (“elongated cell phenotype” in idr0001-A) to form the Set1 histone methyltransferase (HMT). These examples demonstrate that these individual hits are probably not due to off-target effects or characteristics of individual biological models but arise through conserved, specific functions of large macromolecular complexes. This shows the utility and importance of combining phenotypic data of studies from different organisms and scales, and of integrating metadata from independent studies, to generate added value that can enhance the understanding of biological mechanisms and lead to new mechanistic hypothesis and predictions.

The integration of experimental, image and analytic metadata also provides an opportunity to include new functionalities for more advanced visualization and analytics of imaging data and metadata, bringing further added value to the original studies and datasets. As an example, we have added the data analytics tool Mineotaur ^34^ to one of IDR’s datasets (http://idr-demo.openmicroscopy.org/mineotaur/). This allows visual querying and analysis of quantitative feature data. For instance, having shown that components of the Set1 HMT function in controlling cell morphology in *S. pombe* and human cells, we noticed that genes like ASH2L were in the “elongated cell” network based on human cell data (idr0012-A) but not *S. pombe* data, where *ash2*, the *S. pombe* ASH2L orthologue, was not annotated as a cell elongation “hit”. We first noted that *ash2* has a microtubule cytoskeleton phenotype (http://idr-demo.openmicroscopy.org/webclient/?show=well-592371). We then queried the criteria previously used for cell shape hits in the Sysgro screen (idr0001-A) and found that *ash2* fell just below the cutoff originally used in this study to define phenotypic hits for cell shape (Supplemental Information). When combined with results on ASH2L from HeLa cells (idr0012-A, idr0008-B) (Figure 2B) these results suggest that the Set1 HMT has a strongly conserved role in controlling cell shape and the cytoskeleton in unicellular and multicellular organisms.

**Figure 2.**
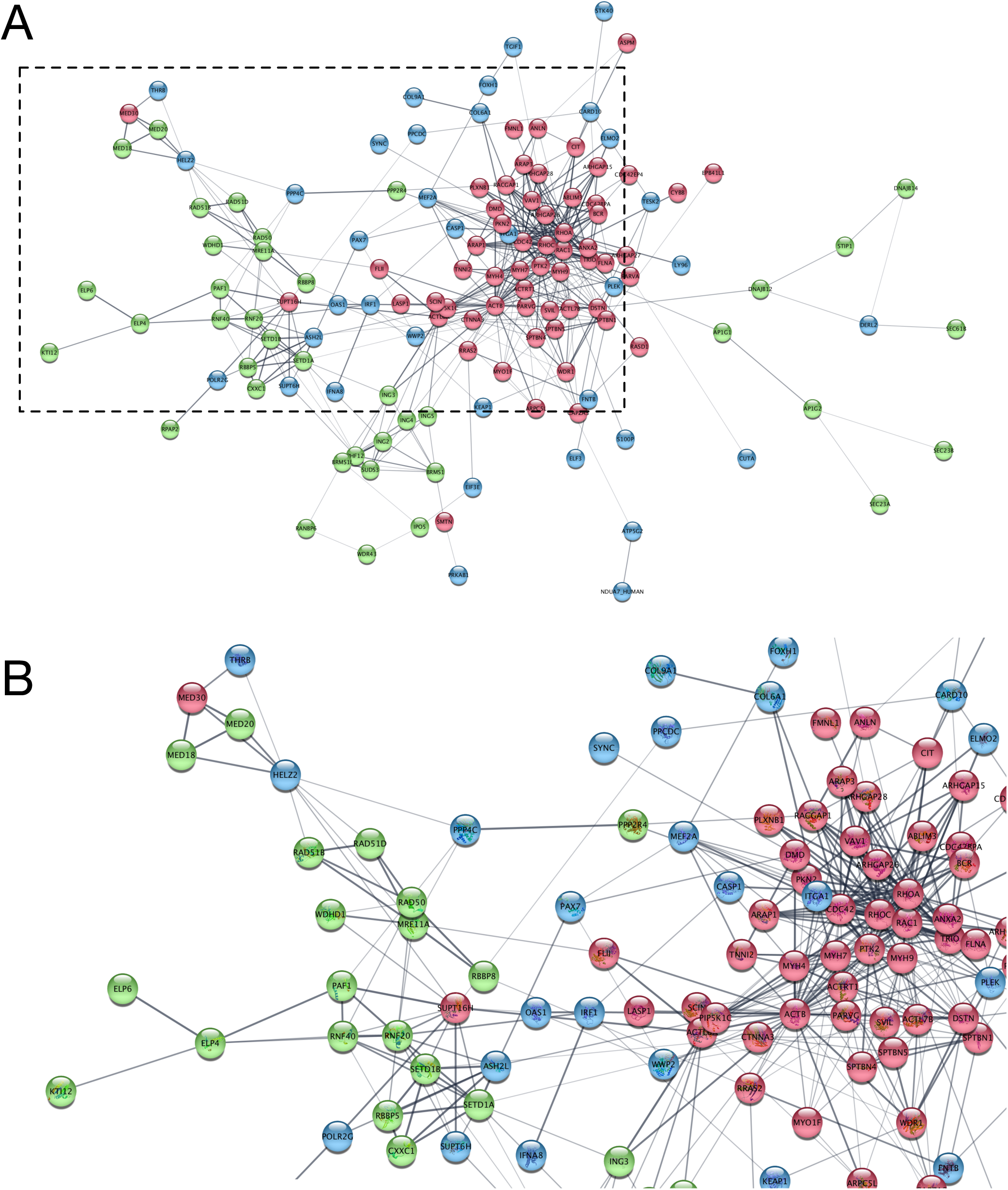
Network Analysis of Genes Linked to the Elongated Cell Phenotype in the IDR. A. Protein-protein interaction network produced in StringDB ^57^ and visualized using Cytoscape (http://www.cytoscape.org/ ^33^) based on the genes linked to the elongated cell phenotype (CMPO_000077) in three distinct studies in IDR. Genes from *S. pombe* (green, idr0001-A, ^5^), HeLa cell morphology (blue, idr0012-A, ^46^) and HeLa Actinome (red, idr0008-B, ^44^) are displayed with linkages (gray) from StringDB. To enable comparisons in Cytoscape, the human orthologues of *S. pombe* genes are used for the genes identified in idr0001-A. It is likely that in the future IDR will receive more submissions that include related phenotypic annotations, so Figure 2 shows one view of an network, based on an evolving set of data. IDR study idr0028 (http://idr-demo.openmicroscopy.org/webclient/?show=screen-1502) also contains experiments annotated with CMPO_0000077/elongated cell phenotype. However this study used targeted libraries, so when included in the network, shows no novel functional interactions. Therefore, data from idr0028 have been omitted from this network, but can be viewed and queried interactively (http://idr-demo.openmicroscopy.org/mapr/phenotype/?value=CMPO0000077) or accessed using the IDR computational resource (https://idr-demo.openmicroscopy.org/jupyter). B. Zoomed view of network in A, with gene names. See Supplemental Information for the list of gene names used in the figure.

### Data Integration and Access

Like most modern on-line resources IDR makes data available through a web user-interface as well as a web-based JSON API. This encourages third-parties to make use of IDR in their own sites. For example, image data in IDR has been linked to study data in BioStudies, thereby extending the linkage of study and image metadata (e.g.,https://www.ebi.ac.uk/biostudies/studies/S-EPMC4704494), and to PhenoImageShare ^35^, an on-line phenotypic repository (e.g.,http://www.phenoimageshare.org/search/?term=&hostName=Image+Data+Repository+(IDR)). These are examples of use of IDR as a service that delivers data for other applications to integrate and reuse.

To add further value and extend the possibilities for reuse of IDR data, we have initiated the calculation of comprehensive sets of feature vectors of IDR image data. For this purpose, we have used WND-CHARM, an open source tool that calculates a broad set of image features ^36^. To date full WND-CHARM features have been calculated for images in idr0002-A, idr0005-A, idr0008-B, idr0009-A, idr0009-B, idr0012-A, and parts of idr0013-A and idr0013-B. Feature calculations for other IDR datasets are in progress. Features are stored in IDR using OMERO’s HDF5-based tabular data store and available through the OMERO API in IDR’s computational resource (see Supplemental Information).

The integration of image-based study phenotypes and calculated features makes IDR an attractive candidate for computational re-analysis. However, given the size of IDR, downloading the full complement of data it contains is impractical. We have therefore built two methods of accessing IDR data. In the first, we have connected IDR to a computational resource that provides remote, API-based access to IDR datasets. This resource authenticates against GitHub and is based on IPython notebooks. This provides a flexible, web browser-based analysis capability for IDR. To demonstrate the utility of this resource, and exemplify its use, we have developed and deposited notebooks that provide visualisation of single tile WND-CHARM features using PCA, access to images annotated with CMPO phenotypes, calculation of gene networks, and calculation of WND-CHARM features for individual images. In particular, we provide notebooks to build interactive versions of Figures 1 and 2 directly from the IDR. Users can access their own analysis notebooks stored in GitHub (https://github.com/IDR/idr-notebooks). IDR’s computational resource is available at https://idr-demo.openmicroscopy.org/jupyter.

In addition, to enable local access to IDR metadata, we have built scripts in Ansible that automate the deployment of the IDR software stack and have made all the databases, metadata and thumbnails in IDR available for download (see Supplemental Information). The downloadable IDR misses the original image data, but contains all image thumbnails and all experimental, phenotypic and analytic metadata associated with IDR images. These can be deployed via the IDR software stack and re-used in a local context.

## Discussion

Making data public and available is a critical part of the scientific enterprise ^37^ (https://wellcome.ac.uk/what-we-do/our-work/expert-advisory-group-data-access) (https://royalsociety.org/topics-policy/projects/science-public-enterprise/report/). To take the next step in facilitating the reuse and meta-analysis of image datasets we have built IDR, a next-generation data technology that integrates and publishes image data and metadata from a wide range of imaging modalities and scales in a consistent format. IDR integrates experimental, imaging, phenotypic and analytic metadata from several independent studies into a single resource, allowing new modes of biological Big Data querying and analysis. As more datasets are added to and integrated with IDR, they will potentiate and catalyze the generation of new biological hypotheses and discoveries.

In IDR, we have linked image metadata from several independent studies. Experimental, imaging phenotypic and analytic metadata are recorded in a consistent format. Rather than declaring and attempting to enforce a strict imaging data standard, IDR provides tools for supporting community formats and releases these as a framework that facilitates data reuse. We hope that the availability of this framework will provide incentives for others to structure their metadata in shareable formats that can be read into IDR or other applications, whether based on OMERO or not. In the future, we can imagine that these and other capabilities could be extended in IDR - or similar repositories that link to IDR - to enable systematic integration, visualization and analytics across imaging studies, thereby helping to harness and capitalize on the exponentially increasing amounts of bio-imaging data that the community generates.

As of this writing, IDR has published 35 reference image datasets grouped into 24 studies (Table 1) and, utilising EMBL-EBI’s Embassy Cloud, has capacity to receive and publish many more. Authors of scientific publications that are already published or under submission can submit accompanying image datasets for publication in IDR, using the metadata specifications and formats we have built. These datasets can be integrated as described above. Once published, the datasets can be browsed and viewed through IDR’s WUI, or queried and re-analysed using the IDR computational resource. Details about the submission process are available under http://idr-demo.openmicroscopy.org/about/submission.html.

Moreover, IDR software and technology is open source, so it can be accessed and built into other image data publication systems. This supports the building of technology and installations that integrate and publish bio-image data for the scientific community, allowing discoveries and predictions similar to what we have shown in Figure 2. IDR therefore functions both as a resource for image data publication and as a technology platform that supports the creation of on-line scientific image databases and services. In the future, those databases and services may ultimately amalgamate to form resources analogous to the genomic resources that are the foundation of much of modern biology.

## Methods

### Architecture and Population of IDR

IDR (http://idr-demo.openmicroscopy.org) was built using open-source OMERO ^18^ and Bio-Formats ^24^ as a foundation. Deployments are managed by Ansible playbooks along with re-usable roles on an OpenStack-based cloud contained within the EMBL-EBI Embassy resource. Datasets (see Table 1) were collected by shipped USB-drive or transferred by Aspera. Included datasets were selected according to the criteria defined by the Euro-BioImaging/Elixir Data Strategy concept of “reference images” (http://www.eurobioimaging.eu/content-news/euro-bioimaging-elixir-image-data-strategy), which states that image datasets for publication should be related to published studies, linked as much as possible to other resources and be candidates for re-use, re-analysis, and/or integration with other studies.

Experimental and analytic metadata were submitted in either spreadsheets (CSV, XLS), PDF or HDF5 files or a MySQL database, each using its own custom format. We converted these custom formats to a consistent tabular format inspired by the MAGE and ISA-TAB specifications ^22, 23^ and combined into a single CSV file using a custom script (available in https://github.com/IDR/idr-metadata) and imported into OMERO. Imaging metadata and binary data were imported into OMERO using Bio-Formats. Experimental and analytic metadata were stored using OMERO.tables, an HDF5-backed tabular data store used by OMERO. For each dataset, metadata that were valuable for querying and search were copied to OMERO’s key-value-based Map Annotation facility ^38^. This means that different metadata types and elements can be accessed using different parts of the OMERO API, depending on the search and querying capabilities they require. For more information on the construction of queries, see Supplemental Information.

## Acknowledgements

We thank all the study authors who submitted datasets for inclusion in the IDR for their contributions and help in incorporating their datasets. We also thank the systems support team at EMBL-EBI, in particular Richard Boyce, David Ocana, Charles Short, and Andy Cafferkey for their support of the project’s use of the Embassy Cloud. We are particularly grateful to Simon Jupp for help with adding and defining new ontology terms. The IDR project was funded by the BBSRC (BB/M018423/1) and Horizon 2020 Framework Programme of the European Union under grant agreement No. 688945 (Euro-BioImaging Prep Phase II). Updates to OMERO and Bio-Formats were supported by the Wellcome Trust (095931/Z/11/Z) and Horizon 2020 Framework Programme of the European Union under grant agreement No. 634107 (MULTIMOT). RECS was funded by a BBSRC Responsive Mode grant (BB/K006320/1), a European Research Council Starting Researcher Investigator Grant (SYSGRO) and the University of Bristol.

## References

1. Uhlen, M. et al. Proteomics. Tissue-based map of the human proteome. Science 347, 1260419 (2015).

2. Hawrylycz, M.J. et al. An anatomically comprehensive atlas of the adult human brain transcriptome. Nature 489, 391–399 (2012).

3. Armit, C. et al. eMouseAtlas, EMAGE, and the spatial dimension of the transcriptome. Mammalian genome: official journal of the International Mammalian Genome Society 23, 514–524 (2012).

4. Neumann, B. et al. Phenotypic profiling of the human genome by time-lapse microscopy reveals cell division genes. Nature 464, 721–727 (2010).

5. Graml, V. et al. A genomic Multiprocess survey of machineries that control and link cell shape, microtubule organization, and cell-cycle progression. Dev Cell 31, 227–239

6. Koh, J.L. et al. CYCLoPs: A Comprehensive Database Constructed from Automated Analysis of Protein Abundance and Subcellular Localization Patterns in Saccharomyces cerevisiae. G3 (Bethesda) 5, 1223–1232 (2015).

7. Gonczy, P. et al. Functional genomic analysis of cell division in C. elegans using RNAi of genes on chromosome III. Nature 408, 331–336. (2000).

8. Fowlkes, C.C. et al. A quantitative spatiotemporal atlas of gene expression in the Drosophila blastoderm. Cell 133, 364–374 (2008).

9. Lawson, C.L. et al. EMDataBank.org: unified data resource for CryoEM. Nucleic Acids Res. 39, D456–464 (2011).

10. Hill, E. Announcing the JCB DataViewer, a browser-based application for viewing original image files. J. Cell Biol. 183, 969–970 (2008).

11. Orloff, D.N., Iwasa, J.H., Martone, M.E., Ellisman, M.H. & Kane, C.M. The cell: an image library-CCDB: a curated repository of microscopy data. Nucleic Acids Res 41, D1241–1250 (2013).

12. Iudin, A., Korir, P.K., Salavert-Torres, J., Kleywegt, G.J. & Patwardhan, A. EMPIAR: a public archive for raw electron microscopy image data. Nat Methods 13, 387–388 (2016)

13. McEntyre, J., Sarkans, U. & Brazma, A. The BioStudies database. Molecular systems biology 11, 847 (2015).

14. Petryszak, R. et al. Expression Atlas update--an integrated database of gene and protein expression in humans, animals and plants. Nucleic Acids Res 44, D746–752 (2016).

15. UniProt, C. UniProt: a hub for protein information. Nucleic Acids Res 43, D204–212

16. Yates, A. et al. Ensembl 2016. Nucleic Acids Res 44, D710–716 (2016).

17. Jupp, S. et al. The cellular microscopy phenotype ontology. Journal of biomedical semantics 7, 28 (2016).

18. Allan, C. et al. OMERO: flexible, model-driven data management for experimental biology. Nat Methods 9, 245–253 (2012).

19. Gorgolewski, K.J. et al. NeuroVault.org: a web-based repository for collecting and sharing unthresholded statistical maps of the human brain. Frontiers in neuroinformatics 9, 8 (2015).

20. Masuzzo, P. et al. An open data ecosystem for cell migration research. Trends Cell Biol 25, 55–58 (2015).

21. Brazma, A., Krestyaninova, M. & Sarkans, U. Standards for systems biology. Nat Rev Genet 7, 593–605 (2006).

22. Sansone, S.A. et al. Toward interoperable bioscience data. Nat Genet 44, 121–126 (2012).

23. Rayner, T.F. et al. A simple spreadsheet-based, MIAME-supportive format for microarray data: MAGE-TAB. BMC Bioinformatics 7, 489 (2006).

24. Linkert, M. et al. Metadata matters: access to image data in the real world. J. Cell. Biol. 189, 777–782 (2010).

25. Goldberg, I.G. et al. The Open Microscopy Environment (OME) Data Model and XML File: Open Tools for Informatics and Quantitative Analysis in Biological Imaging. Genome Biol. 6, R47 (2005).

26. Heriche, J.K. et al. Integration of biological data by kernels on graph nodes allows prediction of new genes involved in mitotic chromosome condensation. Mol Biol Cell 25, 2522–2536 (2014).

27. Simpson, J.C. et al. Genome-wide RNAi screening identifies human proteins with a regulatory function in the early secretory pathway. Nat Cell Biol 14, 764–774 (2012).

28. Shah, G.N. et al. Carbonic anhydrase IV and XIV knockout mice: roles of the respective carbonic anhydrases in buffering the extracellular space in brain. Proc Natl Acad Sci U S A 102, 16771–16776 (2005).

29. Scheibe, R.J. et al. Carbonic anhydrases IV and IX: subcellular localization and functional role in mouse skeletal muscle. Am J Physiol Cell Physiol 294, C402–412 (2008).

30. Wandernoth, P.M. et al. Role of carbonic anhydrase IV in the bicarbonate-mediated activation of murine and human sperm. PLoS One 5, e15061 (2010).

31. Rebello, G. et al. Apoptosis-inducing signal sequence mutation in carbonic anhydrase IV identified in patients with the RP17 form of retinitis pigmentosa. Proc Natl Acad Sci U S A 101, 6617–6622 (2004).

32. Yang, Z. et al. Mutant carbonic anhydrase 4 impairs pH regulation and causes retinal photoreceptor degeneration. Hum Mol Genet 14, 255–265 (2005).

33. Cline, M.S. et al. Integration of biological networks and gene expression data using Cytoscape. Nat Protoc 2, 2366–2382 (2007).

34. Antal, B., Chessel, A. & Carazo Salas, R.E. Mineotaur: a tool for high-content microscopy screen sharing and visual analytics. Genome Biol 16, 283 (2015).

35. Adebayo, S. et al. PhenoImageShare: an image annotation and query infrastructure. Journal of biomedical semantics 7, 35 (2016).

36. Orlov, N. et al. WND-CHARM: Multi-purpose image classification using compound image transforms. Pattern Recognition Letters 29, 1684–1693 (2008).

37. Boulton, G., Rawlins, M., Vallance, P. & Walport, M. Science as a public enterprise: the case for open data. Lancet 377, 1633–1635 (2011).

38. Li, S. et al. Metadata management for high content screening in OMERO. Methods (2015).

39. Breker, M., Gymrek, M. & Schuldiner, M. A novel single-cell screening platform reveals proteome plasticity during yeast stress responses. J Cell Biol 200, 839–850 (2013).

40. Thorpe, P.H., Alvaro, D., Lisby, M. & Rothstein, R. Bringing Rad52 foci into focus. J Cell Biol 194, 665–667 (2011).

41. Toret, C.P., D’Ambrosio, M.V., Vale, R.D., Simon, M.A. & Nelson, W.J. A genome-wide screen identifies conserved protein hubs required for cadherin-mediated cell-cell adhesion. J Cell Biol 204, 265–279 (2014).

42. Fong, K.W. et al. Whole-genome screening identifies proteins localized to distinct nuclear bodies. J Cell Biol 203, 149–164 (2013).

43. Srikumar, T. et al. Global analysis of SUMO chain function reveals multiple roles in chromatin regulation. J Cell Biol 201, 145–163 (2013).

44. Rohn, J.L. et al. Comparative RNAi screening identifies a conserved core metazoan actinome by phenotype. J Cell Biol 194, 789–805 (2011).

45. Doil, C. et al. RNF168 binds and amplifies ubiquitin conjugates on damaged chromosomes to allow accumulation of repair proteins. Cell 136, 435–446 (2009).

46. Fuchs, F. et al. Clustering phenotype populations by genome-wide RNAi and multiparametric imaging. Molecular systems biology 6, 370 (2010).

47. Karsenti, E. et al. A holistic approach to marine eco-systems biology. PLoS Biol 9, e1001177 (2011).

48. Wawer, M.J. et al. Toward performance-diverse small-molecule libraries for cell-based phenotypic screening using multiplexed high-dimensional profiling. Proc Natl Acad Sci U S A 111, 10911–10916 (2014).

49. Breinig, M., Klein, F.A., Huber, W. & Boutros, M. A chemical-genetic interaction map of small molecules using high-throughput imaging in cancer cells. Molecular systems biology 11, 846 (2015).

50. Sero, J.E. et al. Cell shape and the microenvironment regulate nuclear translocation of NF-kappaB in breast epithelial and tumor cells. Molecular systems biology 11, 790 (2015).

51. Barr, A.R. & Bakal, C. A sensitised RNAi screen reveals a ch-TOG genetic interaction network required for spindle assembly. Sci Rep 5, 10564 (2015).

52. Lawo, S., Hasegan, M., Gupta, G.D. & Pelletier, L. Subdiffraction imaging of centrosomes reveals higher-order organizational features of pericentriolar material. Nat Cell Biol 14, 1148–1158 (2012).

53. Szymborska, A. et al. Nuclear pore scaffold structure analyzed by super-resolution microscopy and particle averaging. Science 341, 655–658 (2013).

54. Dickerson, D. et al. High resolution imaging reveals heterogeneity in chromatin states between cells that is not inherited through cell division. BMC Cell Biol 17, 33 (2016).

55. Pascual-Vargas, P. et al. RNAi screens for Rho GTPase regulators of cell shape and YAP/TAZ localisation in triple negative breast cancer. Sci Data 4, 170018 (2017).

56. Yang, W. et al. Regulation of Meristem Morphogenesis by Cell Wall Synthases in Arabidopsis. Curr Biol 26, 1404–1415 (2016).

57. Szklarczyk, D. et al. STRING v10: protein-protein interaction networks, integrated over the tree of life. Nucleic Acids Res 43, D447–452 (2015).

